## Supplementary Information for "An all-in-one pipeline for the *in vitro* discovery and *in vivo* testing of *Plasmodium falciparum* malaria transmission blocking drugs"

**THIS SUPPLEMENTARY MATERIALS FILE INCLUDES**

Materials and Methods

Figs. S1 to S8

Table S1 to S2

Data files S1 to S5

### Materials and Methods

**Workflow for automated image analysis of Hoechst-/MitoTracker-stained stage V gametocytes.**

| Step 1 | Setup | | |
|---|---|---|---|
| | | Image Names: | Channels: |
| | | Hoechst | HOECHST |
| | | MitoTracker | TRITC |

This step defines the channels used for analysis.

| Step 2 | Find Blobs | | |
|---|---|---|---|
| | | Source | MitoTracker |
| | | Approximate Minimum With (µm) | 2 |
| | | Approximate Maximum With (µm) | 20 |
| | | Intensity Above Local Background | 300 |
| | | Result | MitoTracker signal |

This step identifies signals above local background fluorescence in the TRITC channel. Note that the cytoplasm of gametocytes, rather than the mitochondrion only, is included in the "MitoTracker signal" mask (result) using this relatively low intensity threshold. Compare comments in steps 4 and 5.

| Step 3 | Remove Boarder Objects | | |
|---|---|---|---|
| | | Source | MitoTracker signal |
| | | Result | Mitochondria w/o border objects |

This step removes objects situated at the image border of the "MitoTracker signal" mask.

| Step 4 | Filter Mask | | |
|---|---|---|---|
| | | Objects Source | Mitochondria w/o border objects |
| | | Marker Source | Mitochondria w/o border objects |
| | | Measurement | Total Area |
| | | Filter Type | Min Filter |
| | | Maximum Value | 15 |
| | | Include Min/Max Values | No |
| | | Measurement | Ell. Form Factor |
| | | Filter Type | Min Filter |
| | | Maximum Value | 1.8 |
| | | Include Min/Max Values | No |
| | | Result | Mitochondria ell. form factor |

This step removes MitoTracker signal falling below a minimum area and crescent shape (compare comments in step 2) from living cells.

| Step 5 | Find Blobs | | |
|---|---|---|---|
| | | Source | MitoTracker |
| | | Approximate Minimum With (µm) | 1 |
| | | Approximate Maximum With (µm) | 10 |

| | | | |
|---|---|---|---|
| | | Intensity Above Local Background | 2500 |
| | | Result | Mitochondria intensity threshold |

Using a high intensity threshold for MitoTracker signals, this step identifies mitochondria of living gametocytes.

| | | | |
|---|---|---|---|
| Step 6 | Find Round Objects | | |
| | | Source | Hoechst |
| | | Approximate Minimum With (µm) | 0.8 |
| | | Approximate Maximum With (µm) | 3 |
| | | Intensity Above Local Background | 250 |
| | | Result | Round Hoechst objects |

This step identifies parasite-infected red blood cells based on the DNA dye Hoechst.

| | | | |
|---|---|---|---|
| Step 7 | Grow Objects | | |
| | | Objects Source | Round Hoechst objects |
| | | Grow by | 8 |
| | | Result | Nuclei grown |

This step grows the “Round Hoechst objects” mask by 8 pixels in both dimensions.

| | | | |
|---|---|---|---|
| Step 8 | Remove Border Objects | | |
| | | Objects Source | Nuclei grown |
| | | Result | Nuclei w/o border objects |

This step removes objects situated at the image boarder of the “Nuclei grown” mask.

| | | | |
|---|---|---|---|
| Step 9 | Find Blobs | | |
| | | Source | Hoechst |
| | | Approximate Minimum With (µm) | 100 |
| | | Approximate Maximum With (µm) | 300 |
| | | Intensity Above Local Background | 89 |
| | | Result | Blobs |

This step identifies background signal (large contaminants) in images acquired in the HOECHST channel.

| | | | | |
|---|---|---|---|---|
| Step 10 | Grow Objects | | | |
| | | Objects Source | | Blobs |
| | | Grow by | | 200 |
| | | Result | | Blobs grown |

This step grows the “Blobs” mask by 200 pixels in both dimensions.

| | | | |
|---|---|---|---|
| Step 11 | Remove Marked Objects | | |
| | | Objects Source | Nuclei w/o border objects |
| | | Marker Source | Blobs grown |
| | | Result | nuclei |

This step removes areas with significant background from the “Nuclei w/o border objects” mask.

| | | | |
|---|---|---|---|
| Step 12 | Keep Marked Objects | | |
| | | Objects Source | Mitochondria ell. form factor |
| | | Marker Source | nuclei |
| | | Result | Gametocytes |

This step identifies gametocytes within infected red blood cells.

| | | | |
|---|---|---|---|
| Step 13 | Keep Marked Objects | | |
| | | Objects Source | Gametocytes |
| | | Marker Source | Mitochondria intensity threshold |
| | | Result | Live gametocytes |

This step identifies living gametocytes within the "Gametocytes" mask.

| | | | |
|---|---|---|---|
| Step 14 | Measure Mask | | |
| | Objects to Measure | Mask of Objects | nuclei |
| | | Image to Measure | nuclei |
| | Features within Each Object | Mask of Features | Mitochondria Ell. form factor |
| | | Image to Measure | nuclei |
| | Features within Each Object | Source 1 | Live gametocytes |
| | | Source 2 | Gametocytes |

This step quantifies the number of viable gametocytes among infected red blood cells.

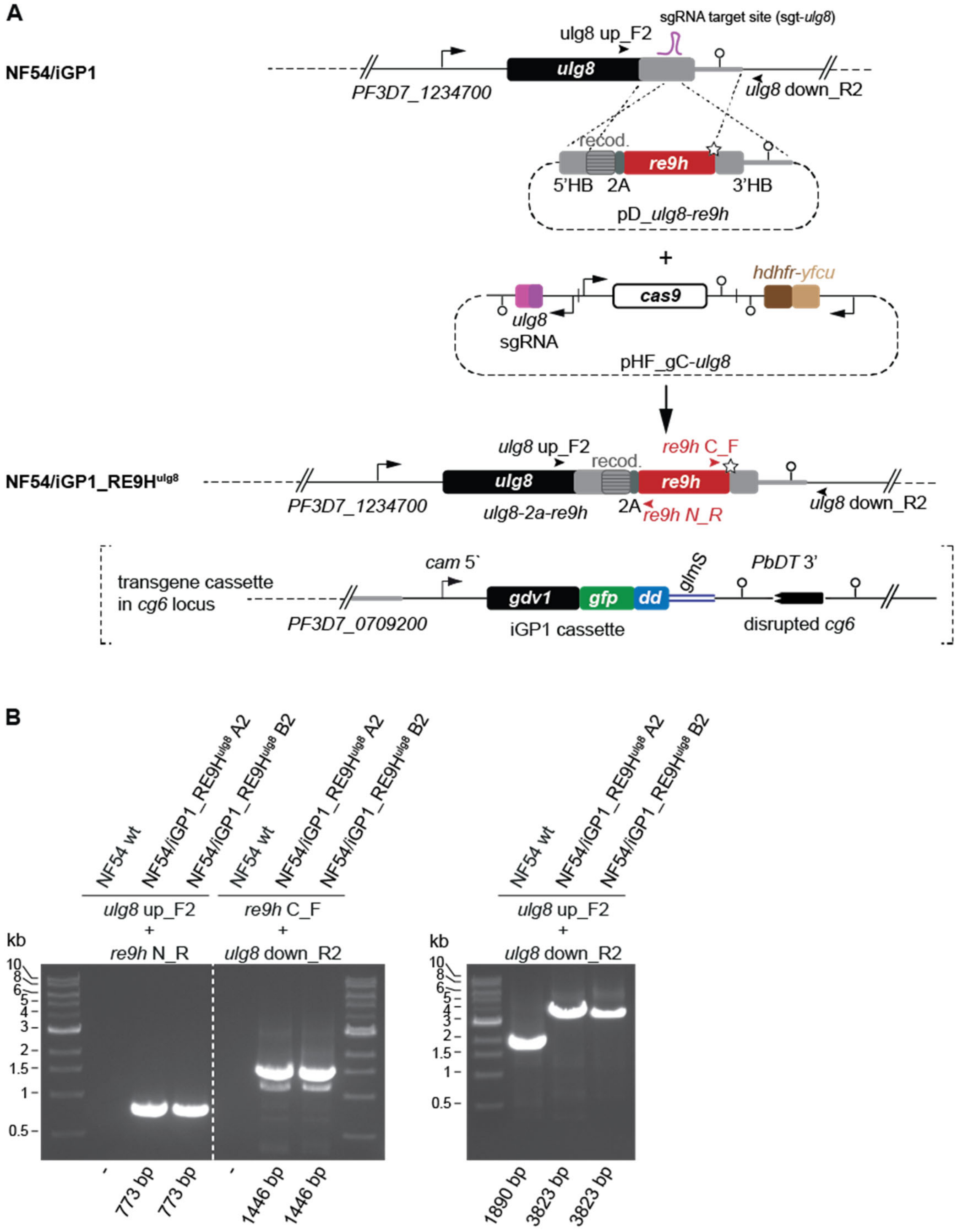

**Fig. S1. Generation of the NF54/iGP1_RE9H$^{ulg8}$ gametocyte reporter line. (A)** Schematic map of the endogenous *ulg8* locus (PF3D7_1234700) (top), the pD_*ulg8_re9h* donor and pHF_gC-*ulg8* plasmids (center), and the gene-edited *ulg8* locus expressing a ULG8-2A-RE9H fusion (bottom) in NF54/iGP1_RE9H$^{ulg8}$ parasites. The relative position of the sgt_*ulg8* sgRNA target sequence is shown in purple. The pD_*ulg8-re9h* donor plasmid contains the 5' and 3' homology boxes (HB) for homology-directed repair (light grey), flanking the re-codonized 3' end of the *ulg8* gene (shaded light grey) followed by sequences encoding the 2A split peptide (dark grey) and the RE9H luciferase (red). The STOP codon is indicated by a white star. The pHF_gC-*ulg8* plasmid contains a SpCas9 expression cassette (white), the sgRNA expression cassette (pink-purple) and the *hdhfr-fcu* positive-negative drug selection marker cassette (brown-grey). Oligonucleotide binding sites used to confirm successful tagging of the *ulg8* gene by PCR on gDNA are indicated by arrowheads. The schematic in brackets at the bottom shows the inducible *gdv1-gfp-dd-glmS* over-expression cassette previously inserted into the non-essential *cg6* locus (PF3D7_0709200) in NF54/iGP1 parasites *(67)*. *cam* 5', *P. falciparum* calmodulin promoter; PbDT 3', *P. berghei dhfr-ts* terminator. glmS, glmS riboswitch element. (**B**) Results of diagnostic PCRs performed on gDNA from NF54 wild-type parasites (control) and two NF54/iGP1_RE9H$^{ulg8}$ clones (A2 and B2). Primer combinations detecting the 5' and 3' recombination events are shown in the left panel. Primer combinations detecting the presence and absence of the *ulg8* wild-type locus in NF54 wild-type and NF54/iGP1_RE9H$^{ulg8}$ parasites, respectively, and the presence of the *ulg8-2a-re9h* fusion gene specifically in NF54/iGP1_RE9H$^{ulg8}$ parasites are shown in the right panel. NF54/iGP1_RE9H$^{ulg8}$ clone B2 has been selected for further experiments.

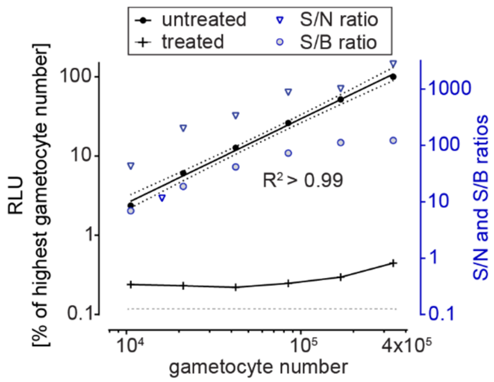

**Fig. S2. Signal-to-noise (S/N) and signal-to-background (S/B) ratios derived from treated and untreated NF54/iGP1_RE9H$^{ulg8}$ stage V gametocytes.** RE9H-catalyzed bioluminescence signals obtained from a dilution series of NF54/iGP1_RE9H$^{ulg8}$ stage V gametocytes (day 12; 3% hematocrit) either left untreated (black dots) or treated with 50 µM MB (black crosses) for 72 hours. Values on the left y-axis represent RLUs (mean of four technical replicates) normalized to the mean signal emitted from the wells containing the highest gametocyte number. The linear regression line (black), coefficient of determination ($R^2$) and s.e.m. (dotted lines) for untreated gametocytes are indicated. Values on the right y-axis represent the S/N ratios (blue triangles) and S/B ratios (blue circles) calculated based on the RLUs obtained from MB-treated and untreated gametocytes. The dashed line at the bottom reflects the normalized RLUs measured from control wells containing uninfected RBCs.

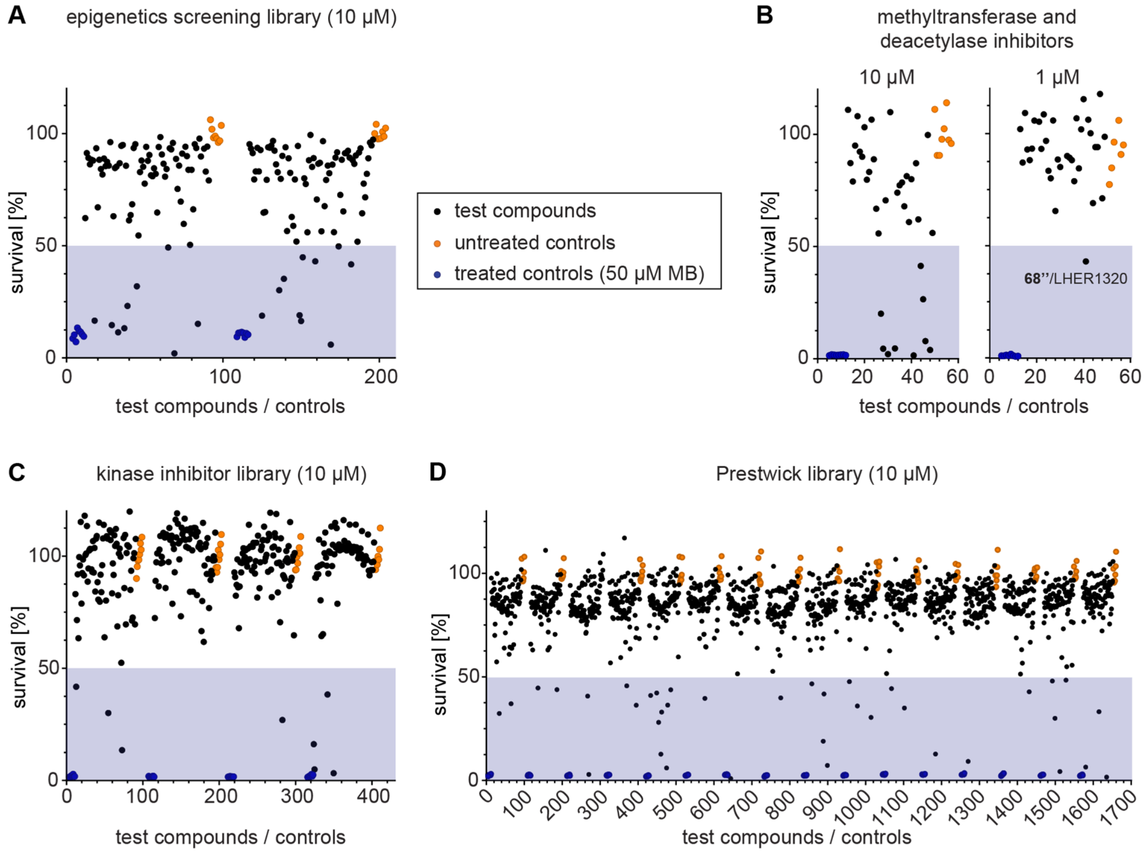

**Fig. S3. Screening of chemical libraries against NF54/iGP1_RE9H$^{ulg8}$ stage V gametocytes.** Results from the primary screening of small- to medium-sized compound libraries. **(A-D)** Effect of compounds of the Epigenetics Screening Library (Cayman Chemical) (10 µM concentration) (A), inhibitors of human DNMTs, HKMTs and histone deacetylases (10 µM and 1 µM concentration) (compound **68''**/LHER1320 with >50% inhibitory activity 1 µM is highlighted), human kinase inhibitors (SelleckChem, Enzo Life Sciences) (10 µM concentration) (C) or compounds of the Prestwick Chemical Library (10 µM concentration) (D) on NF54/iGP1_RE9H$^{ulg8}$ stage V (day 12) gametocyte viability. Each assay plate included eight treated (50 µM MB; blue dots) and untreated (0.1% DMSO; orange dots) samples each as positive and negative controls, respectively. Values on the y-axis represent RLUs normalized to the mean signal emitted from the untreated controls, obtained from a single experiment. Compounds with >50% inhibitory activity are highlighted.

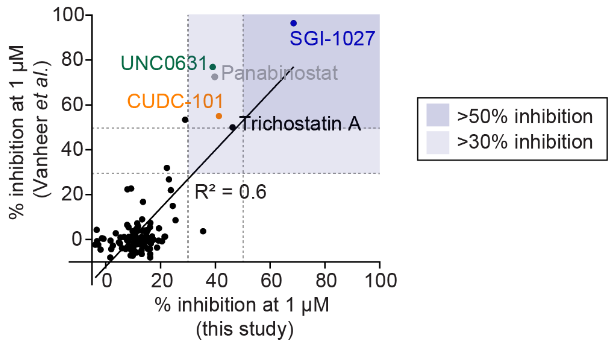

**Fig. S4. Comparison of the gametocytocidal activities of compounds of the Epigenetics Screening Library (Cayman Chemical) screened here and by Vanheer et al. *(63)*.** Scatter plot correlating the inhibitory effects on gametocyte viability of compounds targeting epigenetic enzymes screened at 1 µM concentration against late stage gametocytes (day 10; stage IV/V) in the Vanheer study *(63)* and against mature stage V gametocytes (day 12) in this study. Compounds showing >30% and >50% inhibition are highlighted.

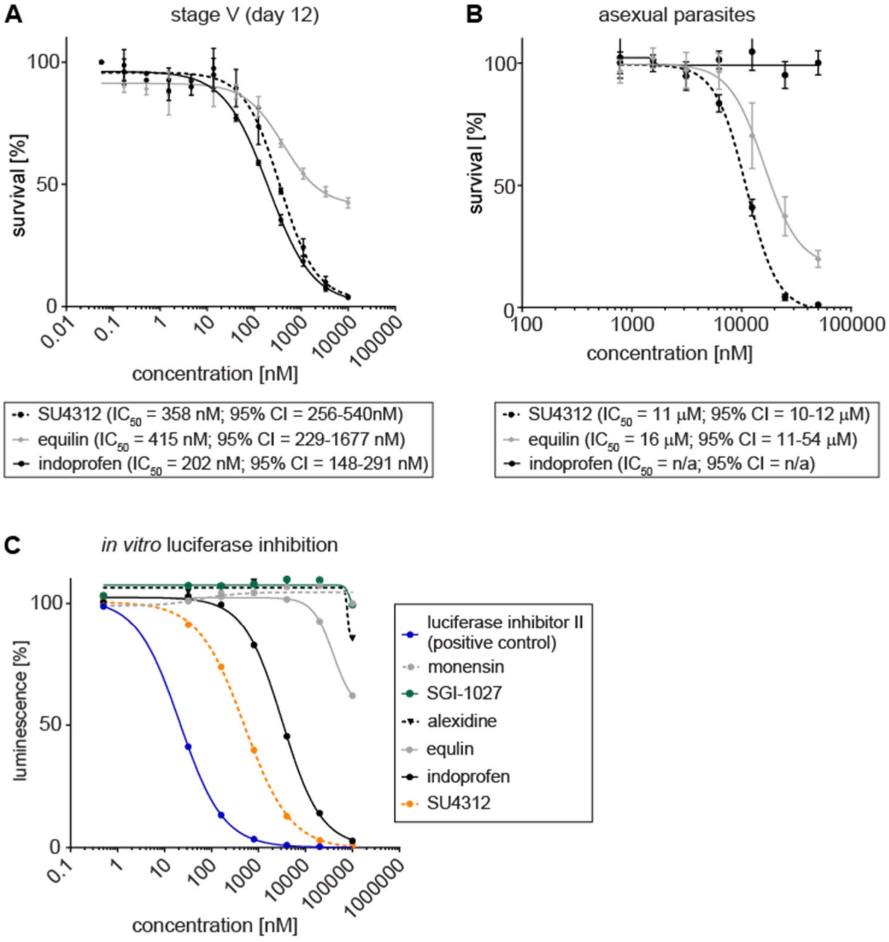

**Fig. S5. Effect of hit compounds against NF54/iGP1_RE9H$^{ulg8}$ stage V gametocytes, asexual parasites and *in vitro* luciferase activity. (A)** Dose-response curves of SU4312, equilin and indoprofen tested against NF54/iGP1_RE9H$^{ulg8}$ stage V gametocytes (day 12). Values on the y-axis represent RLUs normalized to the mean signal emitted from cells exposed to the lowest drug concentration, obtained from three biological replicates (mean ± s.e.m.). **(B)** Dose-response curves of SU4312, equilin and indoprofen tested against NF54 wild-type asexual blood stage parasite multiplication. Values on the y-axis represent [$^3$H]-hypoxanthine incorporation normalized to the mean signal emitted from eight untreated control samples per plate, obtained from three biological replicates (mean ± s.e.m.). $IC_{50}$ values and 95% confidence intervals (CI) are shown below the graphs. **(C)** Dose-response curves of all six hit compounds tested for inhibition of RE9H luciferase activity *in vitro*. Values on the y-axis represent RLUs (mean of two technical replicates) normalized to the mean signal obtained from four untreated control samples. Luciferase inhibitor II was used as a positive control.

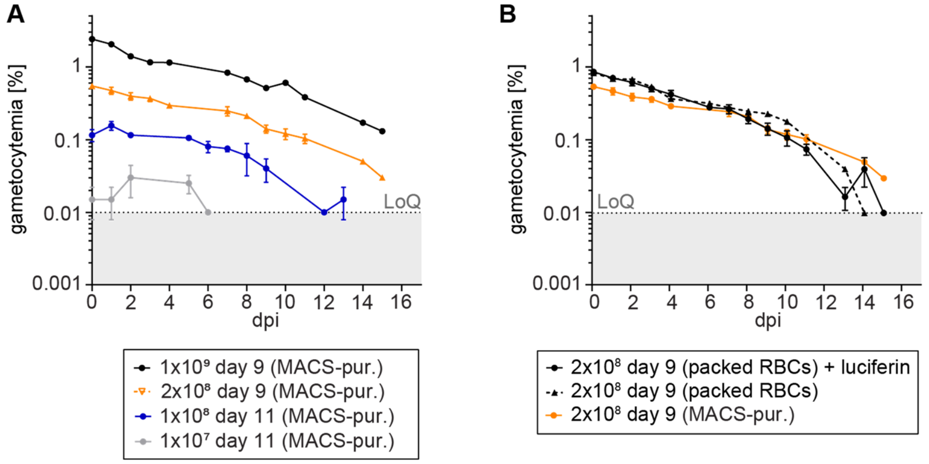

**Fig. S6. Optimisation of the protocol used to infect hRBC-engrafted NODscidIL2R$\gamma^{null}$ mice with NF54/iGP1_RE9H$^{ulg8}$ stage V gametocytes.** Course of peripheral gametocytemia in NF54/iGP1_RE9H$^{ulg8}$-infected mice. **(A)** Mice were infected with 1x10$^7$ (two mice), 1x10$^8$ (two mice), 2x10$^8$ (three mice) or 1x10$^9$ (one mouse) MACS-purified NF54/iGP1_RE9H$^{ulg8}$ gametocytes (day 9 or 11). **(B)** Eight mice were infected with packed RBCs containing 2x10$^8$ NF54/iGP1_RE9H$^{ulg8}$ day 9 gametocytes obtained directly from *in vitro* cultures without prior purification, seven of which received daily injections of 150 mg/kg D-luciferin. Data from mice infected with 2x10$^8$ MACS-purified NF54/iGP1_RE9H$^{ulg8}$ day 9 gametocytes are identical to those shown in panel A. Values on the y-axis represent peripheral gametocytemia (mean ± s.d.) determined by microscopic inspection of Giemsa-stained thin blood smears prepared from tail blood 30 min after infection and once daily thereafter for 15 days. dpi, days post infection; LoQ, limit of quantification.

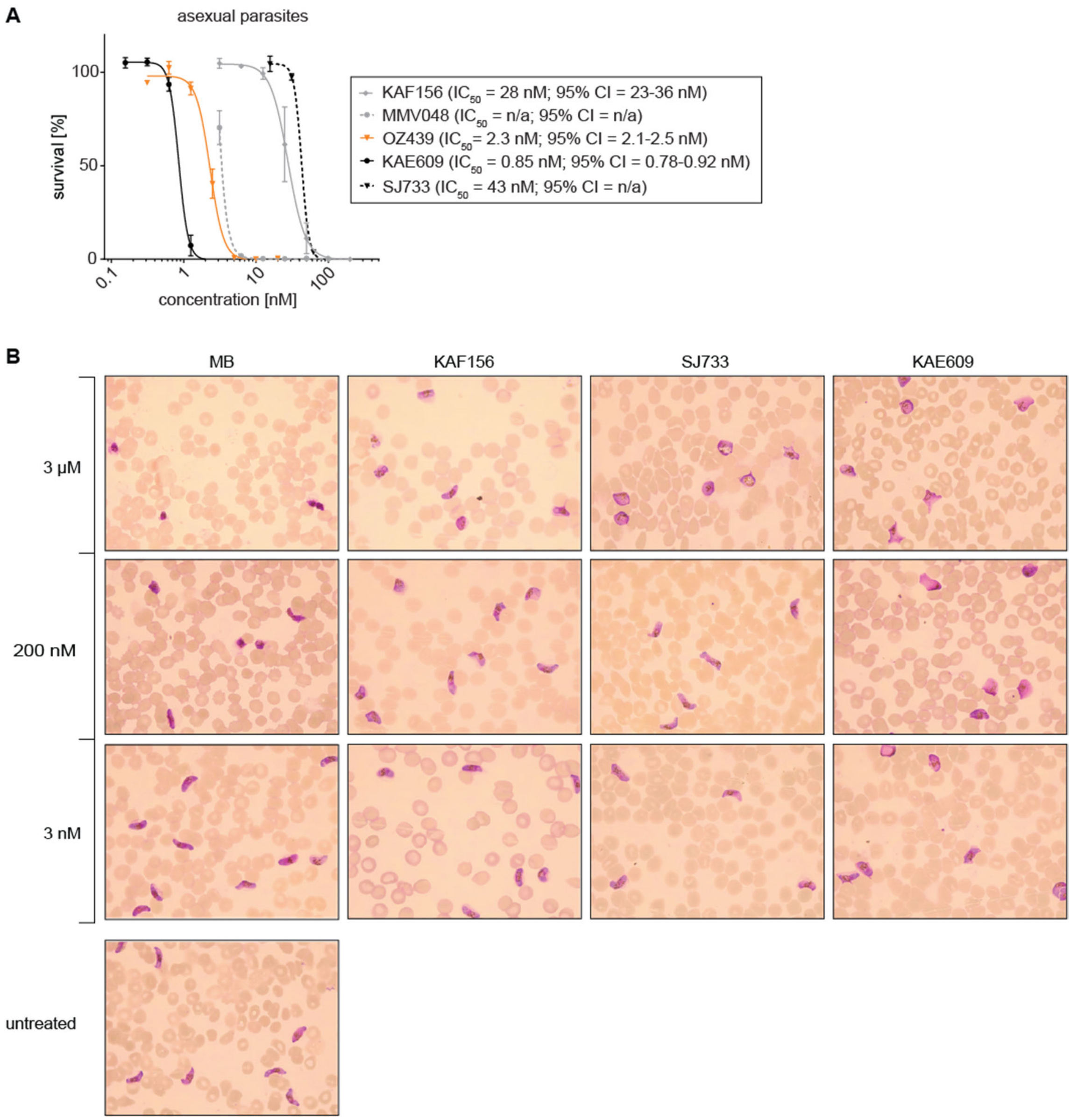

**Fig. S7. Effect of clinical drug candidates on asexual parasite proliferation and stage V gametocyte morphology. (A)** Dose-response curves of clinical drug candidates tested against NF54 wild-type asexual blood stage parasite multiplication. Values on the y-axis represent [$^{3}$H]-hypoxanthine incorporation normalized to the mean signal emitted from eight untreated control samples per plate, obtained from three biological replicates (mean ± s.e.m.). $IC_{50}$ values and 95% confidence intervals (CI) are shown on the right. Note that an $IC_{50}$ value for MMV390048 could not be determined due to lack of data points in the sublethal concentration range. **(B)** Representative images taken from Hemacolor-stained thin blood smears prepared from NF54/iGP1_RE9H$^{ulg8}$ stage V gametocytes (day 12) treated for 72 hours *in vitro* with different concentrations of MB, KAF156/ganaplacide, SJ733 and KAE609/cipargamin. Untreated gametocytes (day 15) are shown as control.

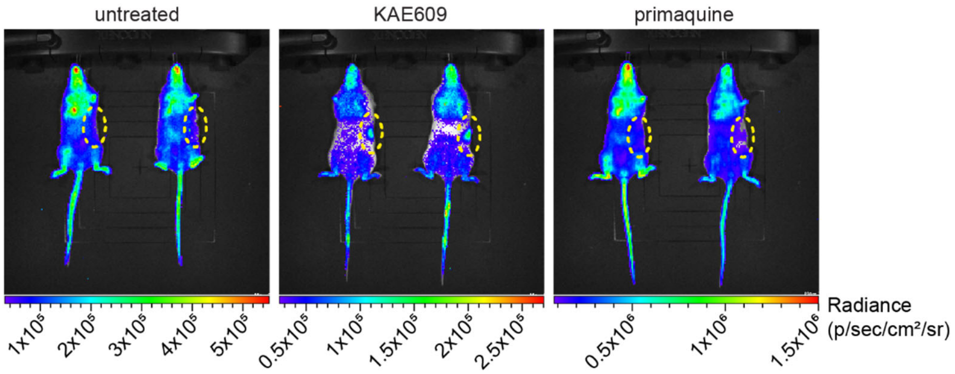

**Fig. S8. Evidence for splenic retention of NF54/iGP1_RE9H$^{ulg8}$ stage V gametocytes in KAE609/cipargamin-treated NSG-PfGAM mice.** Representative ventral images of mice infected with 2x10$^{8}$ NF54/iGP1_RE9H$^{ulg8}$ stage V gametocytes (day 11) and treated one day after infection with 1x40 mg/kg KAE609/cipargamin or 1x50 mg/kg PQ (positive control), or left untreated. Images were recorded after i.v. injection of 150 mg/kg D-luciferin and anesthesia 24 hours post treatment. Pseudocolour heat-maps indicate RE9H-catalysed bioluminescence intensity from low (blue) to high (red). Automated scaling was applied for individual groups to enable comparison of gametocyte densities between heterogeneous groups (e.g. untreated mice with higher gametocyte load compared to low gametocyte load in treated groups). The dashed ovals demarcate the location of the spleen, indicating splenic retention of gametocytes upon treatment with KAE609/cipargamin.

**Table S1. Stage-specific gametocytocidal activity of reference antimalarial drugs and experimental compounds.**

| | $IC_{50}$ [nM (95% CI)] (this study) | | | $IC_{50}$ (nM ± s.d.) *(6)* | | | |
|---|---|---|---|---|---|---|---|
| **Compound** | **III (day 5)** | **IV (day 8)** | **V (day 12)** | **II (day 4)** | **III (day 6)** | **IV (day 8)** | **V (day 12)** |
| Chloroquine | 77 (n/a) | n/a | 41,000 (n/a) | 98 ± 5 | >6,250 | >6,250 | >6,250 |
| Artemisinin | 42 (34-51) | 36 (18-76) | 1,480 (n/a) | 20 ± 10 | 12 ± 2 | 37 ± 3 | >12,500 |
| KDU691 | 861 (699-1,063) | 426 (277-638) | 379 (n/a) | 565 ± 9 | 354 ± 71 | 237 ± 57 | 150 ± 1 |
| Methylene blue | 40 (36-46) | 18 (14-24) | 95 (70-133) | 15 ± 5 | 13 ± 1 | 12 ± 2 | 258 ± 29 |
| Puromycin | 261 (226-301) | 103 (150-371) | 949 (n/a) | 122 ± 45 | 103 ± 39 | 110 ± 38 | 122 ± 48 |

$IC_{50}$ values obtained in this study are derived from three biological replicate dose response assays and 72-hour drug exposure.

**Table S2. Oligonucleotides used for cloning and diagnostic PCRs on gDNA.**

| **Purpose** | **Oligonucleotide** | **Oligonucleotide sequence 5’ → 3’** | **Name of fragment** |
|---|---|---|---|
| pHF_gC-ulg8 | ulg8_sg1F | **tatt**ggtcctttatacgacacagg | sgt_*ulg8* |
| | ulg8_sg1R | **aaac**cctgtgtcgtataaaggacc | |
| pTRIX2-2A-RE9h | 2a_F | *ctgtgcaagagttctagtg*GGTAGCGGAGAAGGAAGAGG | 2a |
| | 2a_R | *ttggcgtcctccatg*ccTGGATTTTCTTCTACATCTCCACAT | |
| pD_ulg8_re9h | 5' box_F | *cgttggccgattcattaatg*CGATGAACATAACCTTGATACA | PCR fragment 1 (5' box) |
| | 5' box_R | *ctccgctaccaTAAAAGAACGCATCC*TTATACT | |
| | 2a-re9h_F | *ggatgcgttcttttatGGTAGCGGAGAAGG*AAGAGGAAGT | PCR fragment 2 (2a-re9h) |
| | 2a-re9h_R | *catatggttgattgcaaTCAGATCTTGCC*CGCCCTTCTTGG | |
| | 3' box_F | *gcaagatctgaTTGCAATCAACCA*TATGACTTAAC | PCR fragment 3 (3' box) |
| | 3' box_R | *cctcttcgctattacgccag*AACCCTAAAGCAAGATGTGTAATA | |
| | PCRA_F | *CTGGCGTAATAGCGAAGAGG* | PCR fragment 4 (pD backbone) |
| | PCRA_R | *CATTAATGAATCGGCCAACG* | |
| PCR on gDNA | ulg8 up_F2 | AGCCTTGTCCTCAAAACTGG | 5' integration |
| | re9h N_R | CTTGATGTTCTTGGCGTCCTCC | |
| | re9h C_F | GAAGGGCGGCAAGATCTG | 3' integration |
| | ulg8 down_R2 | ACATGGACTTACAACCAAAGG | |
| | ulg8 up_F2 | AGCCTTGTCCTCAAAACTGG | wt locus or gene-edited locus |
| | ulg8 down_R2 | ACATGGACTTACAACCAAAGG | |

Names and sequences of oligonucleotides used in this study. Primer binding sites are highlighted in capital letters. Sequences involved in Gibson assembly are highlighted in italics. Single-stranded overhangs required for T4 DNA ligase-dependent cloning of the double-stranded sgRNA target sequence are highlighted in bold font.

**Data file S1.** List of compounds in the Epigenetics Screening Library (Cayman Chemical) and their inhibitory activity against mature stage V gametocytes (day 12) at 10 μM and 1 μM concentrations. RLU, relative luminescence units.

**Data file S2.** Comparison of the inhibitory activities of 101 shared compounds of the Epigenetics Screening Library (Cayman Chemical) screened at 1 μM concentration against mature stage V gametocytes (day 12) in this study and against stage IV/V gametocytes (day 10) by Vanheer et al. *(63)*.

**Data file S3.** List of compounds targeting DNMTs, HKMTs and histone deacetylases and their inhibitory activity against mature stage V gametocytes (day 12) at 10 μM and 1 μM concentrations. RLU, relative luminescence units.

**Data file S4.** List of compounds in the kinase inhibitor library (SelleckChem, Enzo Life Sciences) and their inhibitory activity against mature stage V gametocytes (day 12) at 10 μM and 1 μM concentrations. RLU, relative luminescence units.

**Data file S5.** List of compounds in the Prestwick Chemical Library and their inhibitory activity against mature stage V gametocytes (day 12) at 10 μM and 1 μM concentrations. RLU, relative luminescence units.
